## Supplementary material for "Altered kinetics of circulating progenitor cells in cardiopulmonary bypass (CPB) associated vasoplegic patients: A pilot study": S1_fig

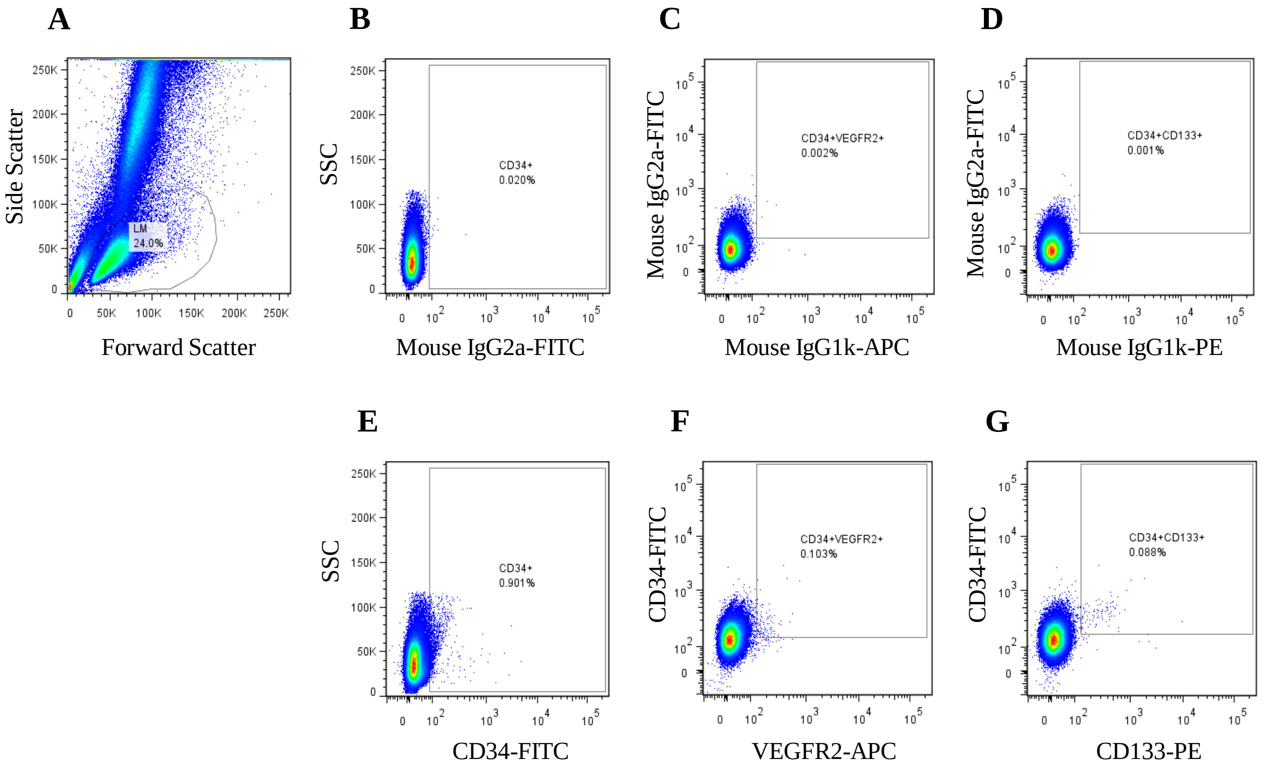


**S1_fig. Gating strategy applied for whole blood.** (A) Morphological gating of the lympho-monocyte fraction using dot plot based on forward and side scatter. (B, C and D) Gating for the CD34-isotype, EPC-isotype, HSC-isotype respectively. (E) Gating for $CD34^+$ cells. (F) Gating for EPCs and (G) Gating for HSCs stained with their respective antibodies.
