## Supplementary material for "Altered kinetics of circulating progenitor cells in cardiopulmonary bypass (CPB) associated vasoplegic patients: A pilot study": S1 Table

**S1 Table. Propensity Matching**

| Patient ID, Group I (no/insignificant vasoplegia) | Euroscore II | Patient ID, Group II (significant vasoplegia) | Euroscore II | Matched |
|---|---|---|---|---|
| 2 | 0.62 | 22 | 0.62 | Yes |
| 6 | 1.9 | 8 | 1.92 | Yes |
| 12 | 1.25 | 25 | 1.62 | Yes |
| 14 | 7.1 | 1 | 6.98 | Yes |
| 15 | 1.8 | 16 | 1.63 | Yes |
| 19 | 2.05 | 4 | 4.27 | No |
| 5 | 0.74 | 9 | 6.07 | No |
| 7 | 0.84 | | | No |
