## Supplementary material for "Altered kinetics of circulating progenitor cells in cardiopulmonary bypass (CPB) associated vasoplegic patients: A pilot study": S2_fig

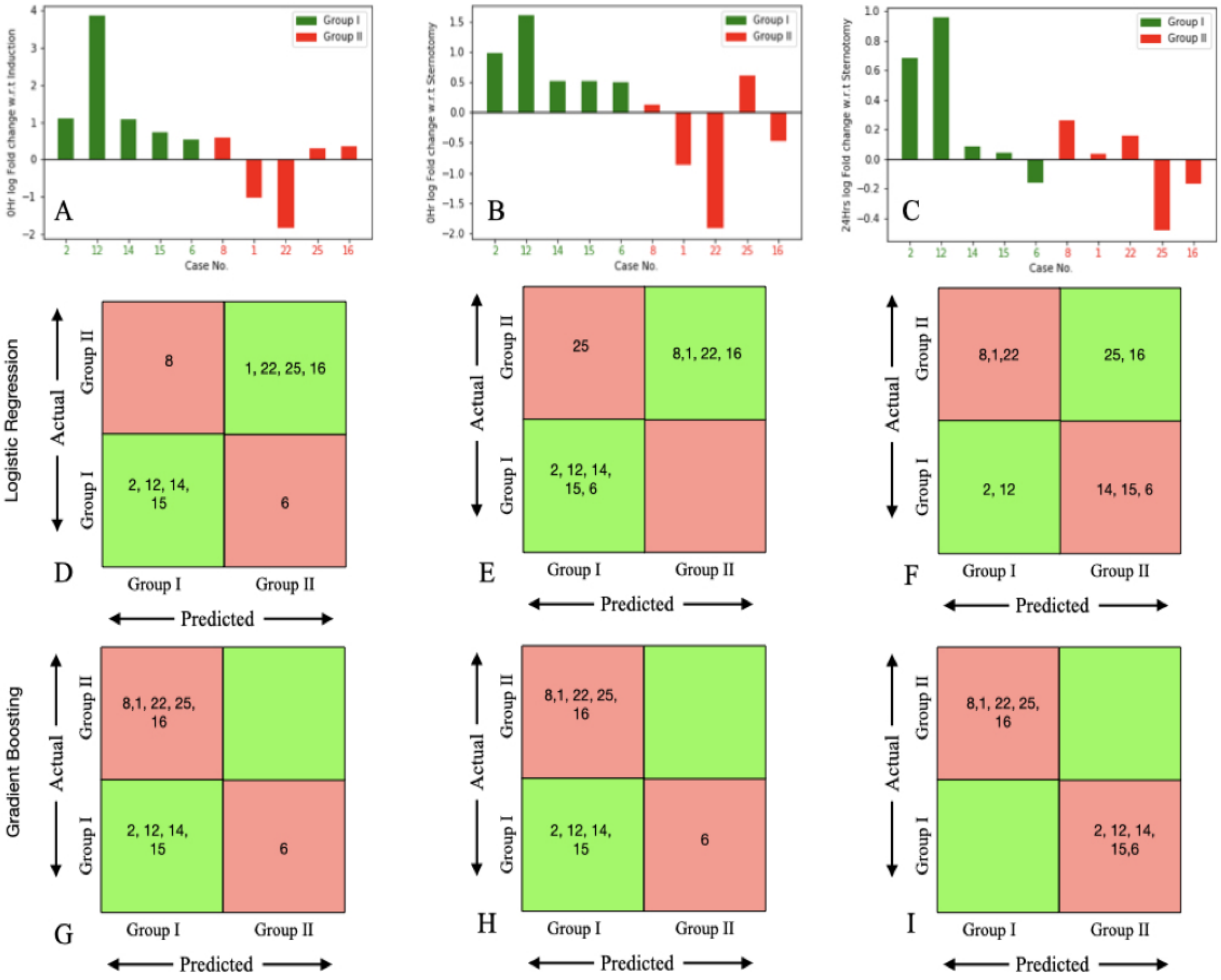


**S2_fig. Propensity matched subgroups of 5 patients under each group (Group I and Group II).** Bar graph showing log fold change of CD34 + marker for each patient, identified by case number, computed at different time points with respect to three baselines: induction, sternotomy and 0 hr. (A) Log fold change at on-pump 0hr with respect to induction. (B) Log fold change for on-pump 0hr time point with respect to sternotomy. (C) Shows 24hr log fold change with respect to sternotomy. Case numbers for patients in G1 (insignificant vasoplegia) are shown in green and those for patients in G2 (significant vasoplegia) are shown in red. (D), (E), (F) show machine learning groupings for G1 and G2 using logistic regression and (G), (H), (I) show the same based on gradient boosting. Correctly classified patients are shown, by case number, in green boxes, and misclassified patients in red boxes
